# Fuzzy-Discernibility Matrix-based Efficient Feature Selection Techniques for Improved Motor-Imagery EEG Signal Classification

**DOI:** 10.1101/2021.03.24.436722

**Authors:** Rajdeep Chatterjee, Debarshi Kumar Sanyal, Ankita Chatterjee

## Abstract

Brain activities, called *brain rhythms*, are the micro-level electrical signals (that is, Electroencephalogram or EEG) generated in our brain while we are performing a task. Even when we imagine a limb movement, it generates the same EEG signals called motor-imagery. Motor-imagery based Brain-computer Interface (BCI) provides a non-muscular means to connect the human brain with limbs through computer-based interpretations. The main aim of this paper is to find a suitable feature-set and a classifier to efficiently classify EEG signals into distinct motor-imagery brain-states. We propose to use sliding temporal window-based approaches for feature extraction from EEG and a mix-bagging classifier which is essentially a bagging-based ensemble of multiple types of learners for motor imagery EEG classification. We observe that mix-bagging with overlapping sliding window-based feature extraction achieves an accuracy of 91.43% on the BCI Competition II Dataset III. To reduce the feature size further, we use a fuzzy discernibility matrix that selects the most discriminative features instead of all the features. This additional feature selection strategy improves the classification accuracy to 92.14% and sets a new state-of-the art result on this dataset.

## 1. Introduction

Electroencephalogram (EEG)-based motor-imagery classification is one of the most extensively explored brain-computer interface (BCI) systems due to its portability and low cost. EEG has its benefits such as portability with high temporal resolution and affordability over other techniques [1, 2, 3, 4]. BCI research is a multi-disciplinary domain that involves subjects such as neuro-science, digital signal analysis, machine intelligence, etc. [5, 6, 7]. Recently, the BCI applications have been developed with hybridized fields such as the Internet of medical things (IoMT), wearable devices and computer-assisted learning and cognition systems [8, 9, 10]. Brain rhythms originate in our brain when we perform any mental task. The brain rhythms are produced even if a person is in a completely relaxed state. In a BCI system, the primary aim is to acquire specific brain patterns and use them (signals) to instruct a computer to perform a particular task. In the classification of these brain patterns, feature extraction algorithms and prediction models play an indispensable role [11, 12]. Activities such as hand movements and foot movements are carried out under the control of the motor cortex region of the brain. Even if we only imagine the limb movements without actually performing them, similar patterns can be observed in the same brain regions (that are activated during the actual limb movements) and thus, they generate similar types of brain rhythms. This imagination-based brain activities is known as motor-imagery (MI) in BCI literature [13, 14]. There are different brain rhythms based on brain functional activities. This paper focuses only on alpha (7 – 13 Hz) and beta (13 – 25 Hz) sub-bands in this paper. Applications such as IoMT and computer-assisted augmented learning work mostly in real-time and deal with *medically significant* life and knowledge-critical data, therefore the solution demands not only high classification accuracy but also lightweight frameworks to obtain it [15, 16, 17, 18].

### 1.1. Related Work

Motor-imagery EEG signal is a non-stationary bio-signal as the spectral (frequency) component varies with time. Different feature extraction techniques have been employed by many researchers for the motor-imagery signal classification problem. In the past [19, 20], wavelet-based feature extraction techniques have been used and compared using a support vector machine (SVM) and multilayer perceptron (MLP) classifiers; the obtained classification accuracies are 85% and 85.71% respectively. Lemm [21] has used Morlet wavelet as features and Bayes quadratic as the classifier to achieve 89.29% accuracy. Similarly, Bashar [22] used multivariate empirical mode decomposition (MEMD) with short-time Fourier transform (STFT) for feature extraction; different variants of classifiers have been used but the best performing one is *K*-Nearest Neighbor (*K*-NN with cosine distance) and obtained 90.71% accuracy. In [23], Wang decomposed the input EEG signals into different frequency sub-bands with second level wavelet packet decomposition technique. The entropy values were computed from the wavelet approximation coefficients and used as a feature-set. The particle swarm optimization technique was used to determine the best discriminating feature subset on which the classification performance of a support vector machine is maximized. Similarly, in [24], Duan used principal component analysis to extract the features and subsequently proposed a decision tree-based optimal feature selection algorithm for the EEG signal classification problem.

Since the early ’90s, multiple studies have been carried out to determine a suitable feature extraction technique for non-stationary bio-signals such as EEG. Schiff et al. [25] introduced Wavelet Transform (WT) as an advantageous feature extraction technique over the then-popular Fourier transforms (FT), Windowed Fourier Transform (WFT) or Short Term Fourier Transform (STFT) [26, 27]. FT captures the spectral, that is, frequency domain information without considering the time domain information (simply, the changes in frequency with variation in time cannot be known). WT is better than WFT due to its no-restriction on the data window size, the length of the window is automatically taken care of in WT. The Heisenberg uncertainty principle states that both the momentum and position of a moving object cannot be observed simultaneously. Similarly, the time and frequency information of a signal at some specific point in the time-frequency space cannot be known. In simple words, what frequency (spectral) component exists at any given time instant cannot be known. The alternative way to overcome this by investigating what frequency (spectral) components are available at any given interval of time instead of a particular point in time. This interval is called a resolution. As STFT keeps a fixed resolution at all times, researchers have shifted from STFT to WT which uses a variable resolution. The computation involved in the Continuous Wavelet Transform (CWT) is not truly continuous, rather it is done based on a finite number of locations. This is a discretized version of the CWT. It is observed from the literature that none of the said techniques, the FT, STFT, and CWT are used in reality for computation due to either it fails to capture the non-stationary characteristics or for their inherent mathematical complexity. The best alternative to avoid this implementation issue as well as to extract the suitable time-frequency information from non-stationary bio-signals is Discrete Wavelet Transform (DWT). In discretized CWT, the obtained wavelet series is nothing but a sampled version of the CWT. The outcome of it is highly redundant in terms of reconstruction of the signal. Thus, it requires a significant amount of computation time as well as resources. On the other hand, (DWT) reduces the redundant information with significantly less computation time while retaining the inherent signal characteristic both in terms of analysis and synthesis of the original signal [28, 29, 30, 31].

Thus, the DWT-based feature extraction technique is well suited for motor-imagery classification over other commonly used feature extraction techniques. Researchers often use *energy and entropy* as features instead of using the *actual wavelet coefficients* for classification [19, 20]. This reduces the feature size. In this paper, we also use the same energy entropy wavelet-based feature extraction technique but in multiple sliding temporal windows, both overlapping and non-overlapping, for a given motor imagery EEG signal. For completeness, we list in Table 1 some of the well-known motor-imagery EEG signal classification techniques used successfully by the researchers.

**Table 1:**
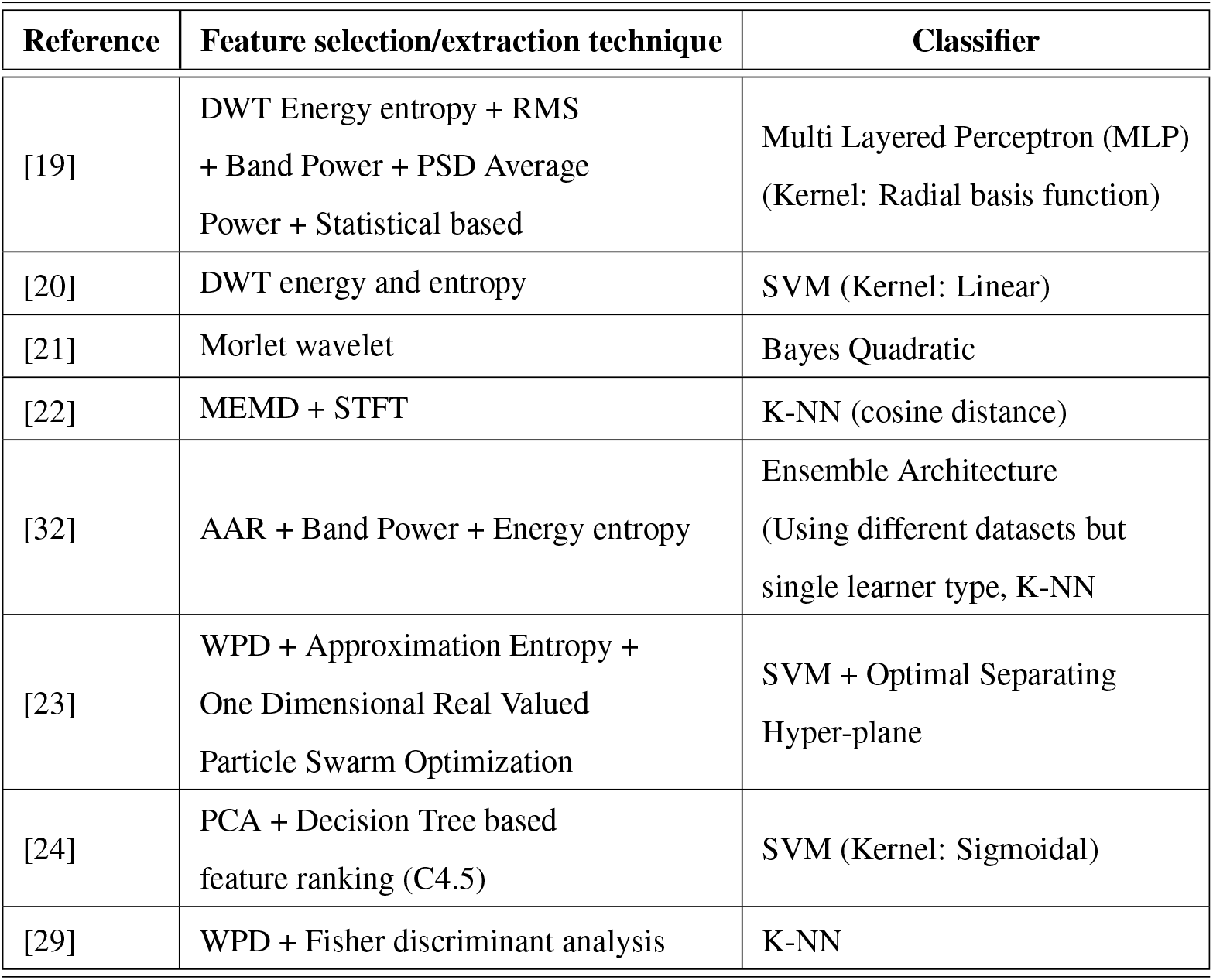
Different best performing techniques used in motor-imagery EEG signal analysis.

| Reference | Feature selection/extraction technique | Classifier |
| --- | --- | --- |
| [19] | DWT Energy entropy + RMS<br>+ Band Power + PSD Average<br>Power + Statistical based | Multi Layered Perceptron (MLP)<br>(Kernel: Radial basis function) |
| [20] | DWT energy and entropy | SVM (Kernel: Linear) |
| [21] | Morlet wavelet | Bayes Quadratic |
| [22] | MEMD + STFT | K-NN (cosine distance) |
| [32] | AAR + Band Power + Energy entropy | Ensemble Architecture<br>(Using different datasets but<br>single learner type, K-NN) |
| [23] | WPD + Approximation Entropy +<br>One Dimensional Real Valued<br>Particle Swarm Optimization | SVM + Optimal Separating<br>Hyper-plane |
| [24] | PCA + Decision Tree based<br>feature ranking (C4.5) | SVM (Kernel: Sigmoidal) |
| [29] | WPD + Fisher discriminant analysis | K-NN |

### 1.2. Motivation

In signal classification problems such as this, one aims to obtain high accuracy. One very important and popular step towards that goal is the use of appropriate feature extraction techniques from signals. Here, the primary focus is to obtain a high level of performance (regarding accuracy) by identifying the most suitable features for classifying the motor-imagery EEG signal.

### 1.3. Contribution

In this paper, we aim to classify motor-imagery EEG signals into two classes: left-hand movement and right-hand movement. Motivated by our prior experience [19, 20, 33], here, we use a DWT-based energy entropy feature extraction technique over multiple – overlapping / non-overlapping – sliding temporal windows on the EEG signal to extract features Then we use the fuzzy discernibility matrix (FDM)-based feature selection technique introduced in our prior work [34] to reduce the feature set size and identify the most discriminating features, i.e., for feature selection. We perform experiments on BCI Competition II Dataset III. With our overlapping DWT-based energy entropy scheme of feature extraction, we obtain classification accuracies of 91.43% and 92.14% *without* feature selection and *with* FDM-based feature selection, respectively. To the best of our knowledge, this is the highest reported accuracy on this dataset. The earlier best result is 90.71% accuracy obtained by Bashar et al. [22].

### 1.4. Organization

The rest of the paper is organized into the following 6 sections. We survey the existing literature in Section 2. The proposed framework is introduced in Section 3 and explained in detail in Section 4. The experimental setup in Section 5. Section 6 presents the results and its analysis. Finally, the study concludes in Section 7.

## 2. Background Concepts

We now elaborate on various concepts that are essential to the methodology we propose in this paper.

### 2.1. Feature extraction techniques

#### 2.1.1. Adaptive Auto-regressive

The *Adaptive Auto-regressive* (AAR) parameters (that is, estimators) represent the characteristics of a signal only for a short period of time. It does not model any trend in the signal. A simple AAR model can be represented by any signal *Y_t_* as [35, 36]:

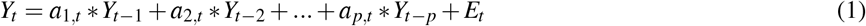

Here, *a*_1*,t*_, ···, *a_or,t_* are AAR estimators in the Eq. 1. *or* indicates the order of the autoregressive model. *E_t_* is the white or random noise. *E_t_* is also known as the prediction error. With smaller error value, the EEG signal is described more accurately by the AAR model.

In this paper, the RLS (Recursive Least Squares) version of AAR models has been used for feature extraction out of 12 available AAR estimations. RLS algorithm is a recursive adaptation of least squares estimators to find the appropriate coefficients that minimize an error estimation related to the input signals. The RLS approach provides an extremely fast convergence for AAR coefficients estimating. As mentioned in [37, 38] that the experimental configurations for the RLS AAR are *a*_1_, *v*_2_ and *UC* = 0. The *a*_1_, *v*_2_ and *UC* indcate the method for updating the co-variance matrix, the method for estimating the innovation variance and the coefficient for model updates, which controls the model adaptation ratio. In our study, two different datasets are generated from the filtered raw EEG signal using AAR feature extraction technique: (1) AAR 12 features dataset, where *or* = 6 for each electrode, and (2) 24 features dataset, where *or* = 12 for each electrode.

#### 2.1.2. Wavelet Energy and Entropy

The *Wavelet energy and entropy* is a well-explored signal processing technique for analyzing non-stationary signals as it well localizes the input signal in time or frequency. *Wavelet transform* seeks to achieve the best trade-off between temporal and frequency resolution. It uses finite basis functions called wavelets. *Wavelet transform* inherently provides the best trade-off between the spectral (frequency) and temporal resolution. It employs the finite basis functions called wavelets. From literature, it is observed that the Discrete Wavelet Transform (DWT)-based energy and entropy feature extraction method is well suited for EEG based motor-imagery classification [19, 20]. DWT is used for feature extraction and its merits have been already discussed in Sub-section 1.1. DWT splits the input signal in half: these are the low-pass and the high-pass sub-bands. The outputs of these sub-bands are also known as approximation coefficients (*A*1) and detail coefficients (*D*1) respectively. One can further decompose the approximation sub-band at multiple levels or scales. The approximation sub-band is again decomposed to another two times to get third-level detail coefficients *D*3 (see Fig. 1) [39]. The third level detail coefficients *D*3 using the Daubechies (db) basis function with filter size 4 is employed to extract features from the input EEG signal [40, 39]. DWT is ideal for downsizing the actual input signal while retaining the property of the original signal with fewer coefficients. It helps in reducing the overall computing process. However, another additional step is used to reduce the dimension of the obtained wavelet coefficients and consolidate the relevant information by implementing Eq. 2 and 3 [19, 40, 41, 42].

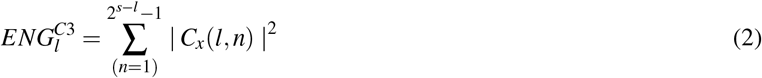

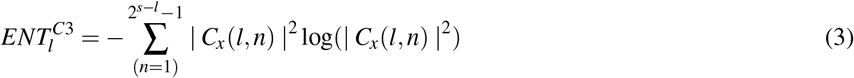

**Figure 1:**
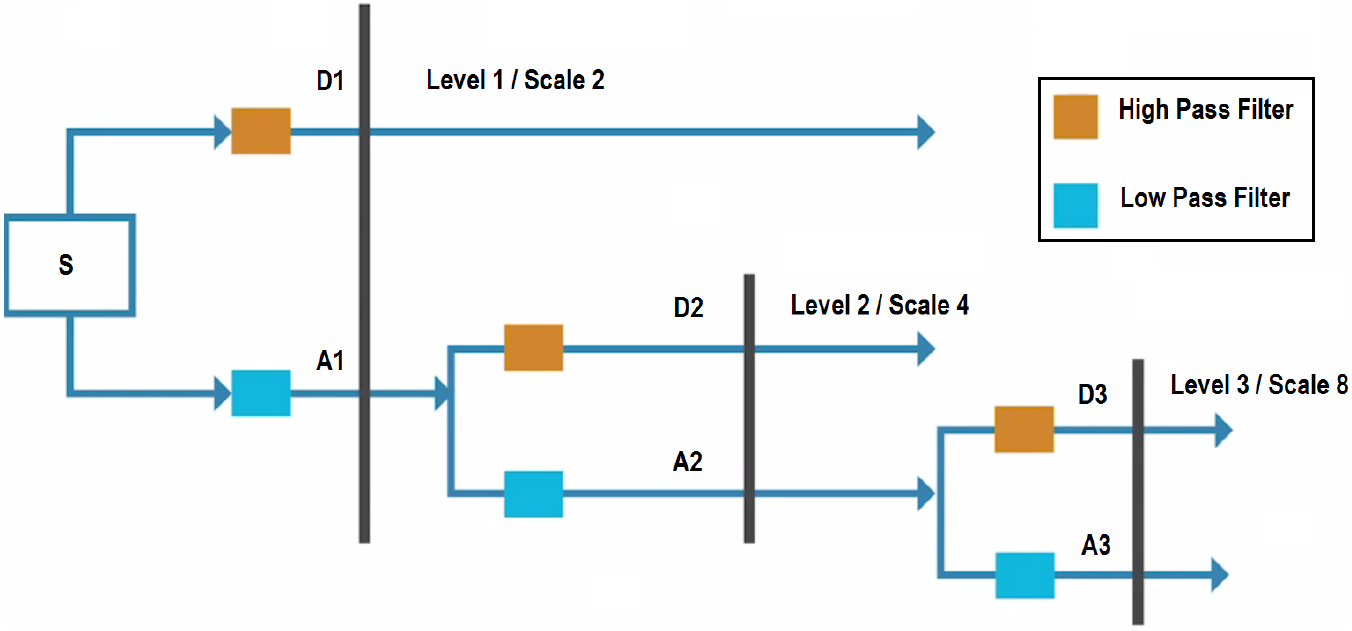
Third level of Discrete Wavelet Transformation into approximation and detail coefficients.

The 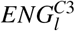 and 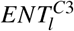 are the computed energy and entropy features for the *l^th^* level of *C*3 electrode. Similarly, energy and entropy for *C*4 electrode are also obtained. *C_x_*(*l, n*) indicates *l^th^* level decomposition and *n^th^* sample for a given input signal.

### 2.2. Feature Selection using Fuzzy Discernibility Matrix

An efficient feature extraction strategy and robust classification technique are important factors for BCI systems. But there is an additional step in between. This step is commonly known as *feature selection*. Different types of univariate and multivariate feature selection techniques are available in the literature. The former [43] are typically fast but provide sub-optimal solutions, whereas the latter [44] are too time-consuming. As the EEG-based decision making needs real-time or near real-time solutions [44, 45], we prefer univariate feature selection techniques for their lightweight computational requirements.

#### 2.2.1. Fuzzy Discernibility Matrix

The fuzzy adaptation of the traditional decision-relative discernibility matrix is called Fuzzy Discernibility Matrix (FDM) and the detail explanation is available in [34, 46]. Earlier, the classical discernibility matrix [47] deals only with the discretized dataset, thus it incurs information loss. The fuzzy variant, that is, FDM has overcome this shortcoming. The proposed algorithm needs a fuzzified dataset as an input instead of the original crisp dataset. Suppose *p* is the total number of features for a given dataset, excluding the decision class. Let the input dataset be denoted by *X^p^* which is a column matrix containing the values from all the trials for the *p^th^* feature. The process begins with the calculations of different means and standard deviations from the actual (non-fuzzified) dataset. The standard deviations (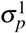 & 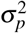) and means (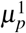 & 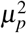) are computed decision class-wise for a feature: left-hand movement (label 1) and right-hand movement (label 2). These are used to generate the corresponding fuzzified sub-features using Eq. 4. The Gaussian membership function is employed to fuzzify each of the features given in Eq. 4 [48]. In Eq. 4, the *σ* and *μ* indicate the standard deviation and the mean of the *p^th^* feature. However, the class labels are kept crisp (that is, binary class labels 1 and 2) in the processed dataset for implementing the decision-relative property of the discernibility matrix.

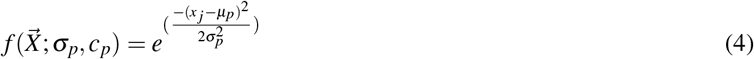

Here, 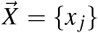 is the non-fuzzy feature value where *j* = {1, 2, …, number of instances}. The *σ_i_* and *μ_i_* are standard deviation and mean of the *i^th^* fuzzy set respectively (due to two decision classes of our used dataset, *i* = 1, 2). We obtain two fuzzy sets from each crisp feature: (i) degree of belongingness to the Left-hand motor-imagery movement for a trial (an instance) based on a *p^th^* feature value; (ii) degree of belongingness to the Right-hand motor-imagery movement for a trial (an instance) based on the same *p^th^* feature value. Therefore, two fuzzy sub-features are generated from each non-fuzzy original feature.

Assume *D_A_* = (*X, A, D*) is a given dataset with Universe *X* = {*x*_1_*, x*_2_*, …, x_m_*} (where *m* indicates the total number of instances in the dataset), conditional attributes (crisp features) *A* and binary decision class *D* = {*L, R*}. After fuzzification, *D_A_* transforms to *F_D_* as explained earlier. In *F_D_*, a crisp feature *c ∈ A* decomposed to 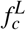 and 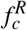 fuzzy values^1^ using Eq. 4. For a set *c*, denote the fuzzified set as *c^*^*. Let *d*(*x_i_*) and *d*(*x_j_*) indicate the decision class types of the *i^th^* and *j^th^* trials respectively.

The proposed fuzzy discernibility matrix of *D_A_* is denoted by *FD_M_*(*F_D_*), where each cell entry is a vector of size *n* = *|A|* defined by fuzzy discern equivalence condition (that is, dissimilarity measure) as follows:

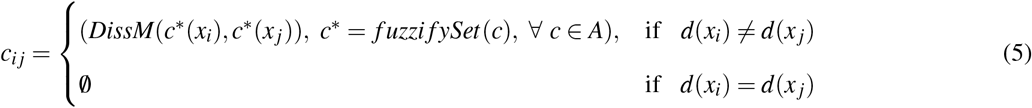

where, *i, j* = 1, 2*, …, m*.

Here, *c_i j_* is a vector of feature-wise dissimilarity values. Each component *t* of the vector *c_i j_* is a a dissimilarity index value that lies between 0 to 1, and discriminates instances *x_i_* and *x_j_* into two decision classes with different degrees of discernibility.

In [46], different dissimilarity functions (*DissM*) have been used to compute feature-wise discrimination between two samples based on the decision classes. Two distinct algorithms have been given in [46]. However, these are rewritten in this paper as: (1) *Algorithm 1* is for computing the Fuzzy Discernibility Matrix, (2) *Algorithm 2* is for identifying the best discriminating feature subset (also known as Reduct). It is observed by the authors through rigorous empirical means that the dissimilarity measure Eq. 6 of this paper (that is, Eq. 10 of [46]) provides the best and consistent feature subset. The dissimilarity measure between two trials using the fuzzy feature-sets 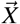 and 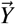 (here in Eq. 6, fuzzy-sets are derived from a crisp feature and binary decision classes set the fuzzy-sets per feature as two) while *i* = 1, …, *N* can be defined as follows:

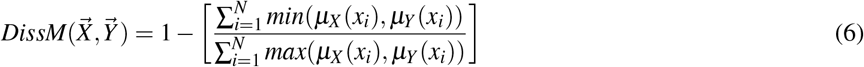

The value of *N* in Eq. 6 suggests the *classNum* (total number of decision classes for a given dataset). The 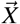 and 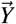 are vectors indicating the fuzzified values *μ_X_* (.) and *μ_Y_* (.) for *x^th^* and *y^th^* samples (*x* ≠ *y*). The dissimilarity function Eq. 3 is derived from established **Fuzzy Similarity Measure** 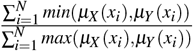 by subtracting it from 1 [49, 50, 51].

The proposed FDM feature selection algorithm can be used on other numerical datasets, but here, it is explained in the context of our study to maintain the relevance. It is a symmetric matrix like its predecessor, only one half of the fuzzy discernibility matrix (excluding the diagonal elements) is computed. After the fuzzification of the crisp dataset, the new feature-set size becomes *ActualFeatureNum × classNum*, where *actualFeatureNum* is the original number of features present in the dataset (after feature extraction) and *classNum* is the number of decision classes.

The feature-wise dissimilarity is calculated for any two instances (that is, trials) iff their class labels are different. It is known as the decision-relative property, and it boosts the information gain for each feature based on discrimination. In Algorithm 1, *t* is the *dissimilarity score*. The dissimilarity value lies between 0 to 1. A value close to 1 suggests full dissimilarity between any two instances for a feature; 0 indicates complete similarity. The increase in dissimilarity value suggests an increase in the discriminating information for a feature.

#### 2.2.2. Reduct: the feature subset

The *Reduct* is a subset of selected features from the original set of features. Each valid entry of fuzzy discernibility matrix (*FD_M_*(*i, j*) termed as *c_i j_* in Eq. 5) is a vector of dissimilarity scores, known as “discernibility vector”. Its length equals to the *ActualFeatureNum*, that is, the number of crisp features of the original dataset. Each vector element indicates a feature-wise discernibility for two different instances. In Algorithm 2, the step 4 sums all the discernibility vectors feature-wise. Then it divides the resultant vector by 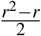 (that is, the number of valid cell entries in the fuzzy discernibility matrix). The normalization is applied at the end to keep the final values within the range of 0 to 1. Thereafter, a sorting mechanism applies to the final *discernibility vector* in descending order. Finally, the user input *s* is used to select the first *s* best discriminating features as *Reduct*.

**Algorithm 1.**
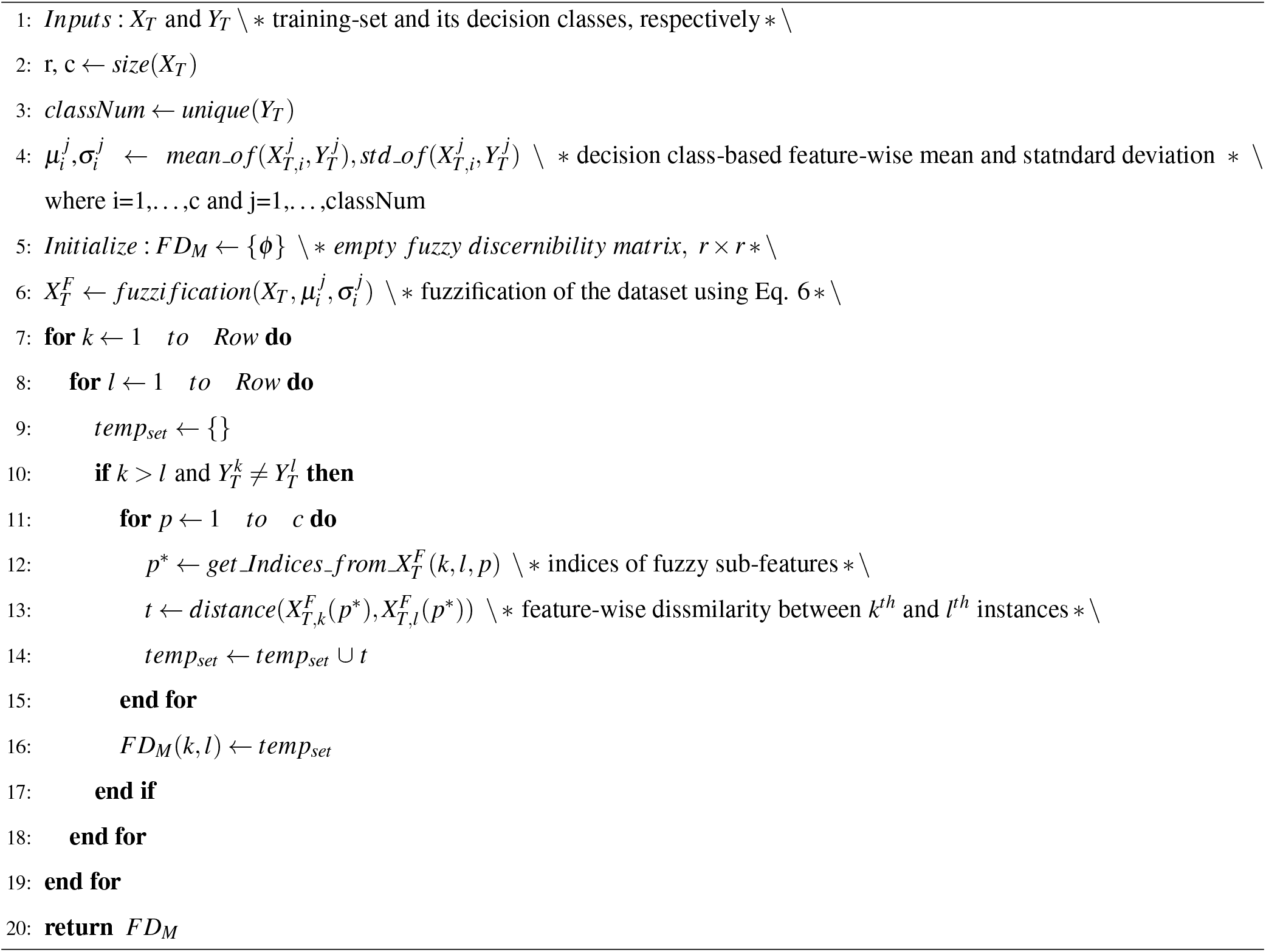
Fuzzy Discernibility Matrix (*FD_M_*)

**Algorithm 2.**
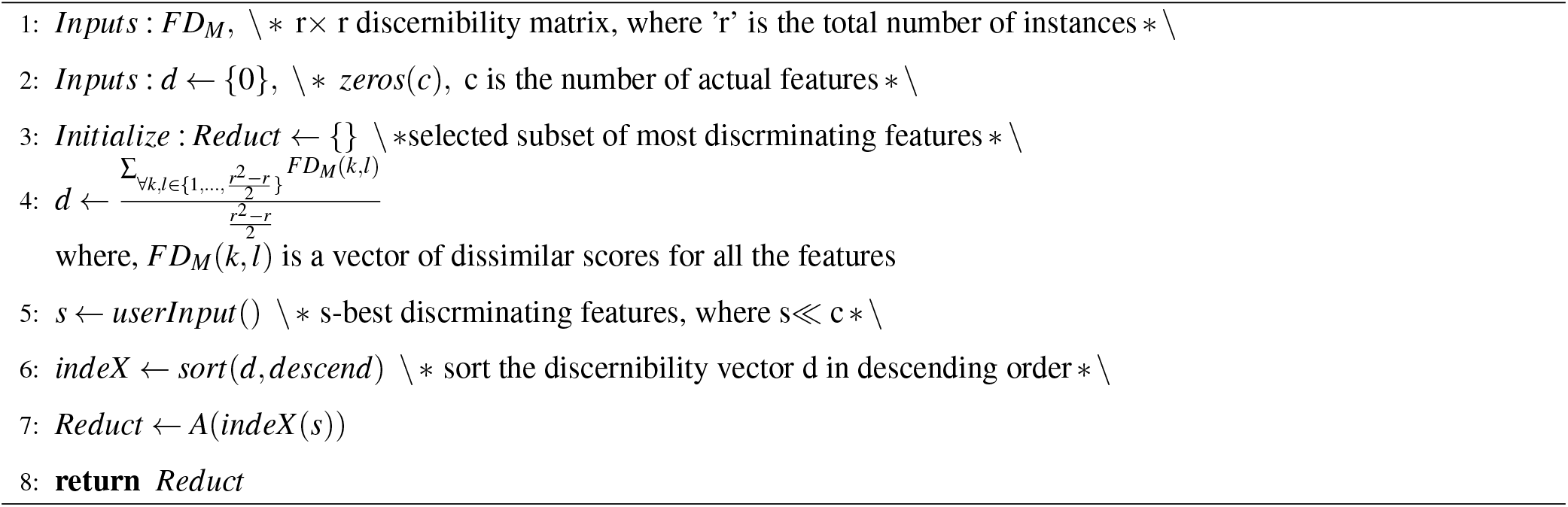
Reduct: feature subset selection

#### 2.2.3. Concept discussion

The real-valued crisp data have been transformed to fuzzy values based on their feature-wise and class-standard-deviations (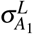 & 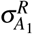) and class-means (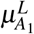 & 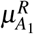). These are calculated for a feature based on those samples which belong to a particular decision class. For example, 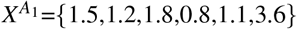 where, the number of samples are 6 and *A*_1_ indicates the feature number. Suppose, the decision class set *D*={R,L,R,L,L,R} (L: Left-hand movement and R: Right-hand movement). Therefore, the values in 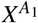 are grouped into two subsets 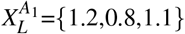 and 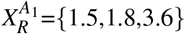 based on two distinct class labels *L* and *R*, respectively. Now, the 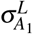 and 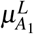 are computed based on sample subset 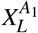. Again, the 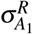 and 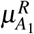 are computed based on sample subset 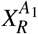.

The red ellipsoid indicates that the operation is performed per feature. The dissimilarity measure Eq. 6 can be used to determine the feature-wise discrimination between any two instances (e.g., instances 1 and 2 are shown in Fig. 2). The green and red markers indicate the fuzzified sub-feature vectors obtained from the actual crisp feature *A*_1_ to be used in computing the dissimilarity in Fig. 3.

**Figure 2:**
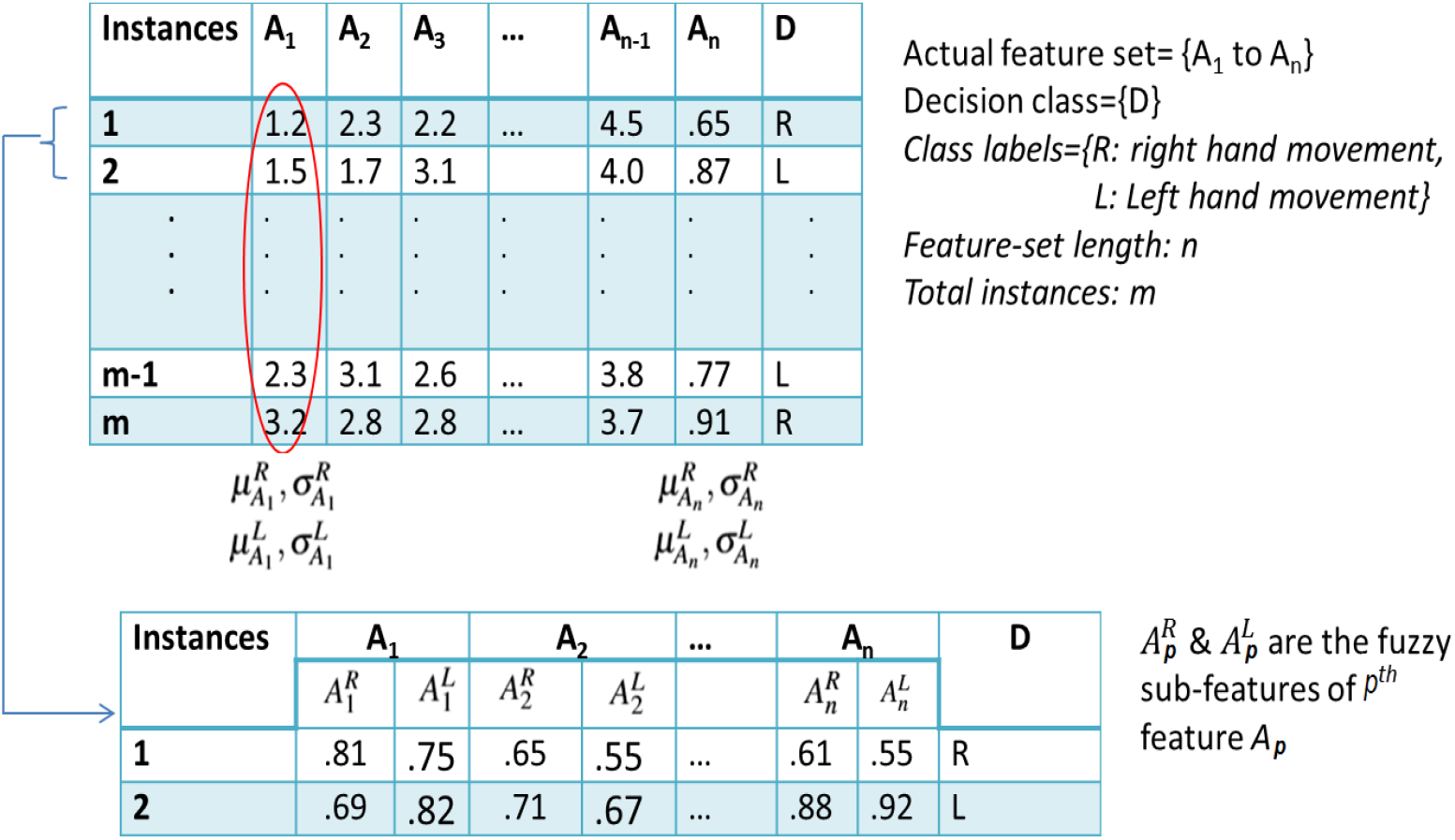
Visual example of the fuzzification step

**Figure 3:**
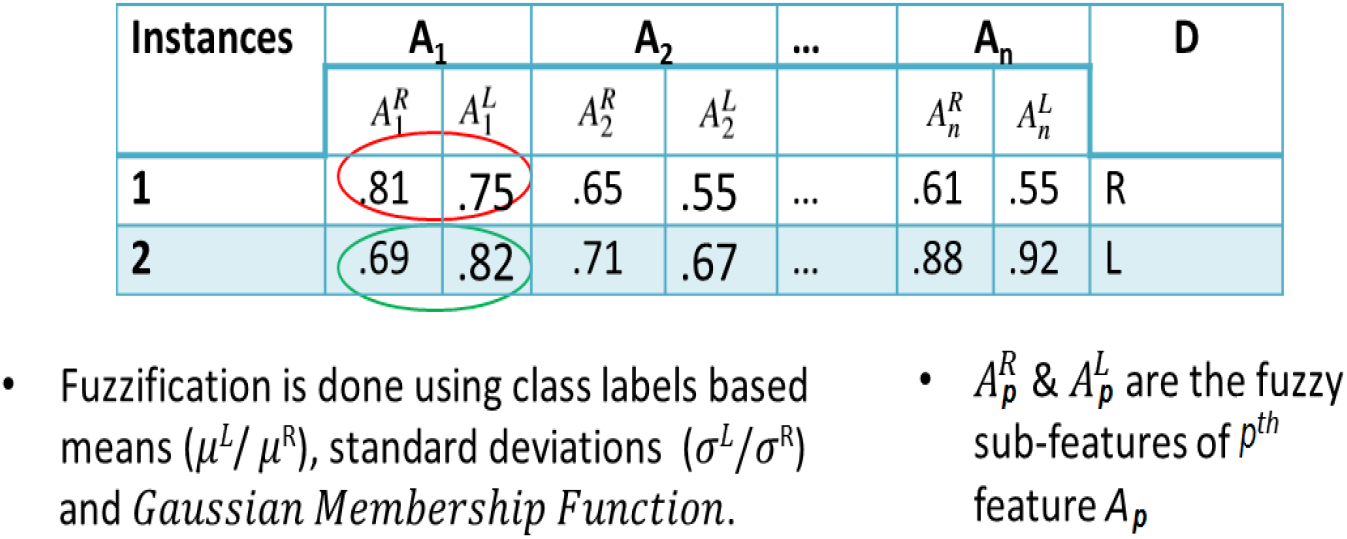
Visual representation of the dissimilarity measure while implementing the Fuzzy Discernibility Matrix

Assume an already fuzzified dataset is given in Table 2 where each feature value indicates a degree of belongingness to different decision classes *R* and *L* for the instances 1 and 2. The fuzzy discernibility matrix obtained from the proposed Algorithm 2 and given in Table 3 for the instances 2 and 1. Each valid entry in the *FD_M_* is a vector of three dissimilarity values. It is so as it has only three crisp features (conditional attributes {*A, B,C*}). In Table 3, The fuzzy discernibility vector between instance 2 and 1 denotes the dissimilarity values: *FD_M_{*2, 1} = [0.88, 0.86, 0.17].

**Table 2:** Example of a Fuzzified Table.

| Instances | A |  | B |  | C |  | D |
| --- | --- | --- | --- | --- | --- | --- | --- |
|  | A<sub>Y</sub> | A<sub>N</sub> | B<sub>Y</sub> | B<sub>N</sub> | C<sub>Y</sub> | C<sub>N</sub> |  |
| 1 | 0.81 | 0.75 | 0.65 | 0.55 | 0.10 | 0.20 | R |
| 2 | 0.69 | 0.82 | 0.71 | 0.67 | 0.88 | 0.92 | L |

**Table 3:** Fuzzy Discernibility Matrix of Table 2.

| Instances | 1 | 2 | 3 | 4 |
| --- | --- | --- | --- | --- |
| 1 |  |  |  |  |
| 2 | [0.88,0.86,0.17] |  |  |  |
| 3 | [1x3] | [1x3] |  |  |
| 4 | ∅ | [1x3] | [1x3] | |

We compute the other valid dissimilarity vectors. The cell entry, for instances 4 and 1 in our example, is ∅, which suggests both the instances belong to the same decision class. Finally, an input *s* is taken from the user to determine the Reduct using Eqs. 7, 8 and 9. The Reduct is the selected set of highest discriminant features *A*(*indeX* (*s*)). (refer Algorithm 2 of [46]).

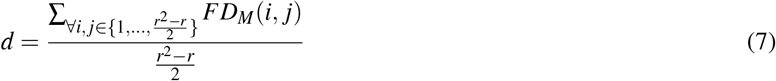

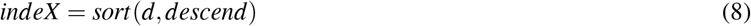

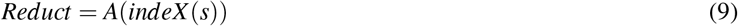

The name of the proposed algorithm is justified as the obtained discernibility matrix uses fuzzy values (degrees of discernibility for all features) instead of direct feature values.

#### 2.2.4. Time Complexity Analysis

Let, the input matrix *X_T_* is of *m × n* dimensions. The number of rows (instances) and the number of columns (attributes) are denoted by *m* and *n* respectively. As per Algorithm 1, three distinct conditional statements are used in steps 7, 8 and 11 (for-loops). The step 10 implements the decision-relative property in Algorithm 1. Due to step 10, the process is also executed only 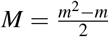 times. Again due to step 11, it is executed for *N* = *n* times for each *M* steps. Therefore, the *Time Complexity* of the proposed algorithm to compute fuzzy discernibility matrix is 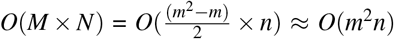 [in Big-Oh asymptotic upper bound notation].

Similarly, in Algorithm 2 the step 6 is executed for 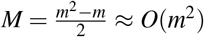. The array needs to be sorted equals the length of *|A|*, that is, (n-1) except the decision class. The step 6 uses Matlab inbuilt “SORT” method which is an quick sort technique, so it gives us *N* = *O*(*nlogn*) as *n* is total number of features. So the final *Time Complexity* for Reduct computation is *O*(*M* + *N*) = *O*(*m*^2^ + *nlogn*) *≈ O*(*m*^2^), where *m > n* [in Big-Oh asymptotic upper bound notation].

### 2.3. Classifiers

Different types of classifiers: Naïve Bayes, *K*-Nearest Neighbor, Support Vector Machine with different kernels, and ensemble classifier (boosting and bagging) have been implemented to compare the performances [52, 53, 54, 55]. The mean of *K*-NN outputs are computed by varying the *K* values from 3 to 35 and the *cosine* distance metric is used for each time. Also, the number of learners in boosting (ENS1 and ENS2) and the number of bags for bagging ensemble approaches (that is ENS3) are averaged by varying them from 10 to 100. It must be mentioned that for all normal ensemble approaches, the learner type is always fixed as *decision tree*. These variations are done to find out the best performing classifier configuration for our experiments.

Besides these classifiers, another classifier named as mixture-bagging (hereafter, *mix-bagging*) ensemble technique is employed in our study borrowed from [56, 57]. The mix-bagging ensemble classifier is formed by using multiple types of weak learners (here, mix-bag-size=5) for different bags. The performance of the mix-bagging classifier that uses *multiple learners* gives the *best* result due to its higher diversity over the single classifier type. Specifically for bagging and mix-bagging, the sample bag sizes are kept at 65% of the actual training dataset (*m_use_*). This configuration of parameters is chosen due to its stable and consistent performance after rigorous trials. Here, the mix-bagging is a modified and improved classical bagging ensemble classifier. It has been executed *iter* times independently. In each iteration, the mix-bagging checks whether the currently obtained accuracy is greater than the previous iteration. If the condition holds *true*, it keeps the latest results and also stores the bags (*indx*, the sub-training-sets for each bag) otherwise it retains the previous result and the ensemble configuration. The details of all the used classifiers are given in Tables 4 and 5.

**Table 4:**
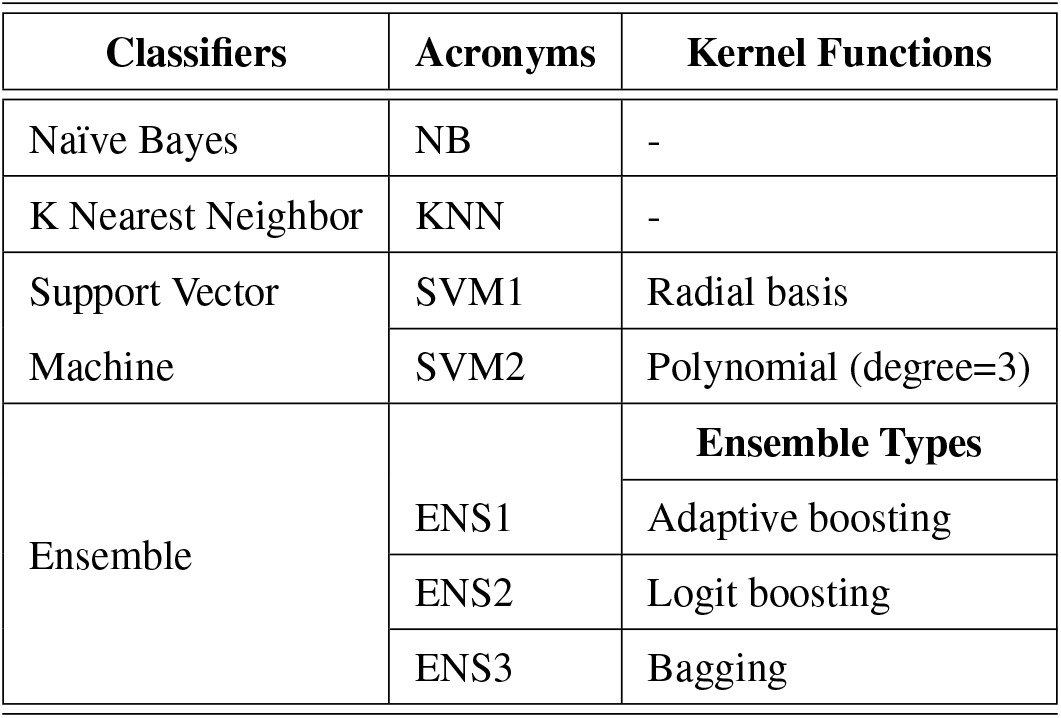
Classifier configurations used in this paper.

**Table 5:** Configuration of the mix-bagging ensemble classifier with 5 different learners.

| Bag # | Learner Types | Parameters |
| --- | --- | --- |
| Bag 1 | Tree | Ovabyclass |
| Bag 2 | KNN (cosine) | K=13 |
| Bag 3 | Discriminant | Linear |
| Bag 4 | Naïve Bayes | - |
| Bag 5 | SVM | Linear |

The majority-voting technique is used for combining the results obtained from multiple predictors of bagging (ENS3) and mix-bagging. It works on the simple principle of voting based on maximum similarity as shown in Fig. 4.

**Figure 4:**
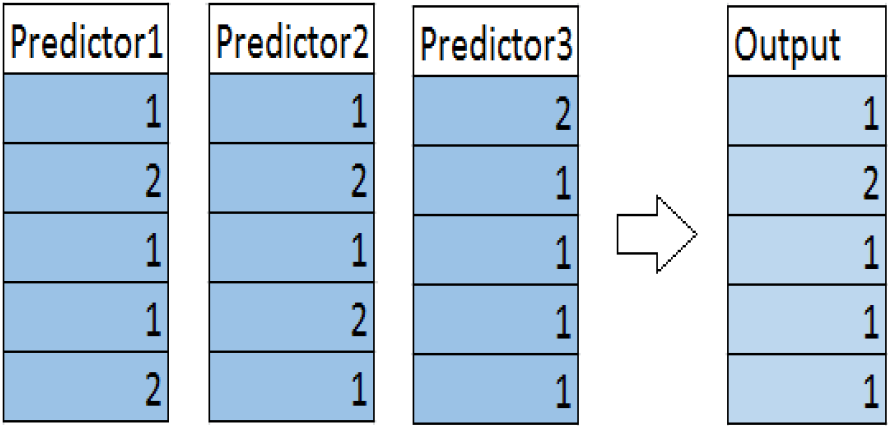
Example of majority-voting with three predicted output vectors

Our proposed mix-bagging classifier brings diversity due to its different types of learners and outperforms the other used alternatives. Even if mix-bagging performs well over other bagging variants with multiple but similar types of learners. In other words, if Support Vector Machine with a linear kernel is used for all the bags instead of using multiple types of classifiers, it under-performs. Similarly, other individual classifiers such as KNN with *K* value 13, NB, LDA and DT classifiers for all the five bags are implemented separately to examine their performance. To validate our claim that mix-bagging introduces more diversity, the *first two features* of the overlapping energy and entropy dataset has been used shown in Fig. 5, that is, Exp-(c) (as it is the best performing dataset in our experiments) already explained in Table **??**.

**Figure 5:**
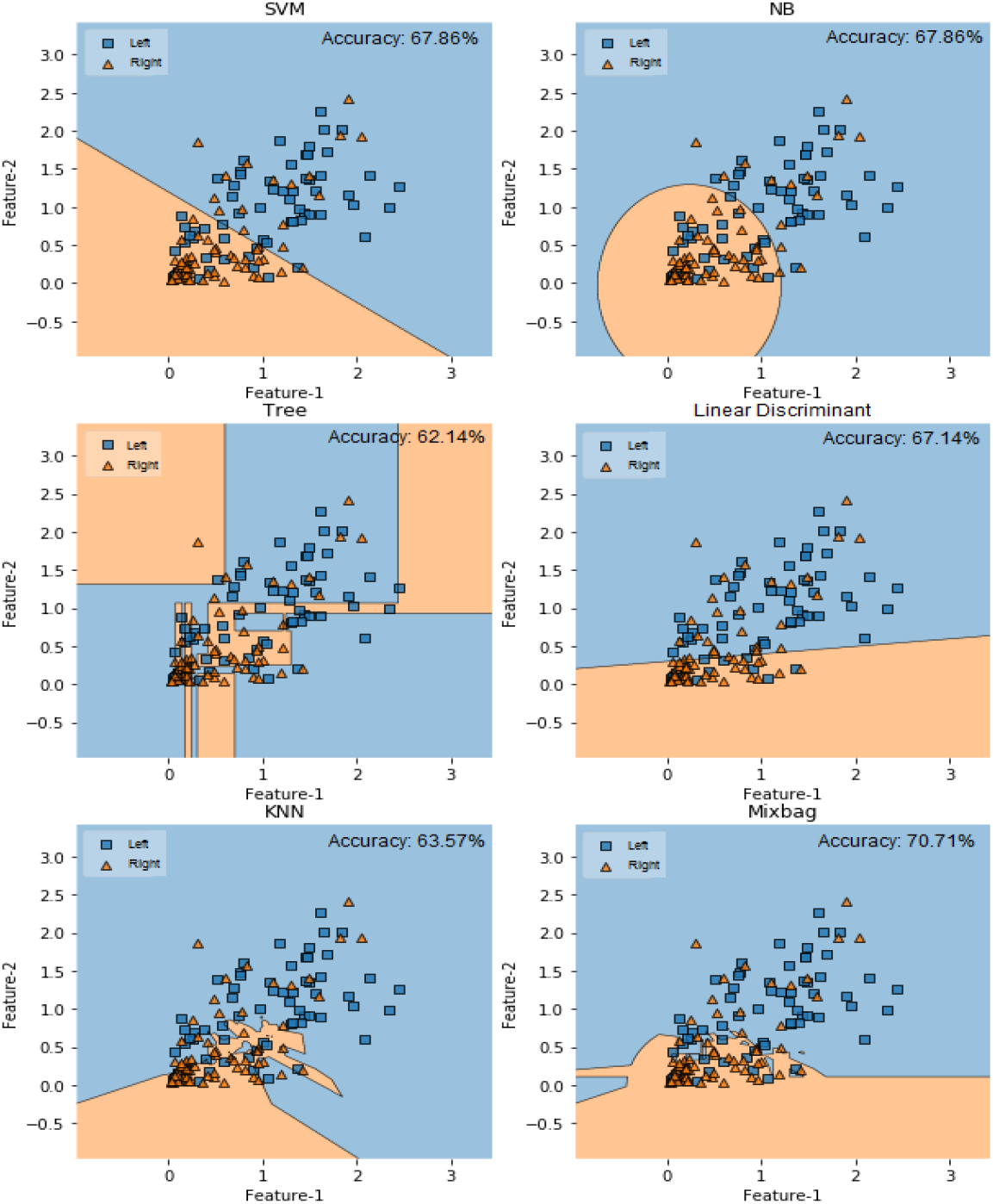
Comparative performance analysis of mix-bag over other alternatives while using first two features of Exp-(c) dataset

It is observed that the proposed mix-bag classifier model works better even with only two features. The decision boundaries for each cases in two dimension feature spaces (Feature-1 *×* Feature-2) are plotted separately. The accuracies mentioned in Fig. 5 are obtained while using the first two features from Exp-(c) dataset (explained in the previous section).

## 3. Overview of the Proposed Framework

Our goal is to introduce an efficient feature selection technique to identify the most discriminating features in the input signals so that we can input this reduced feature set to the classifier. High dimensional feature-space incurs computational overhead in classification. Moreover, the use of all the features does not necessarily assure the best classification accuracy. If some of the features are highly correlated with each other or have minimal discriminatory contribution in the classification model, then it might be better to select the subset of features that are most effective in classification. Feature selection is not only apt for reducing the feature-set size but it also increases the accuracy by discarding redundant non-discriminatory features. Currently, very few such approaches have been implemented for motor-imagery EEG signal classification. It has huge potential in this domain as real-time BCI applications require lightweight prediction models with high accuracy and a small number of features. We aim to employ a newly introduced feature selection technique that can identify the most discriminating features in motor imagery EEG signals and thereby, enhance classification accuracy. Therefore, the visual representation of the proposed framework for an improved EEG-based motor-imagery signal classification is shown in Fig. 6.

**Figure 6:**
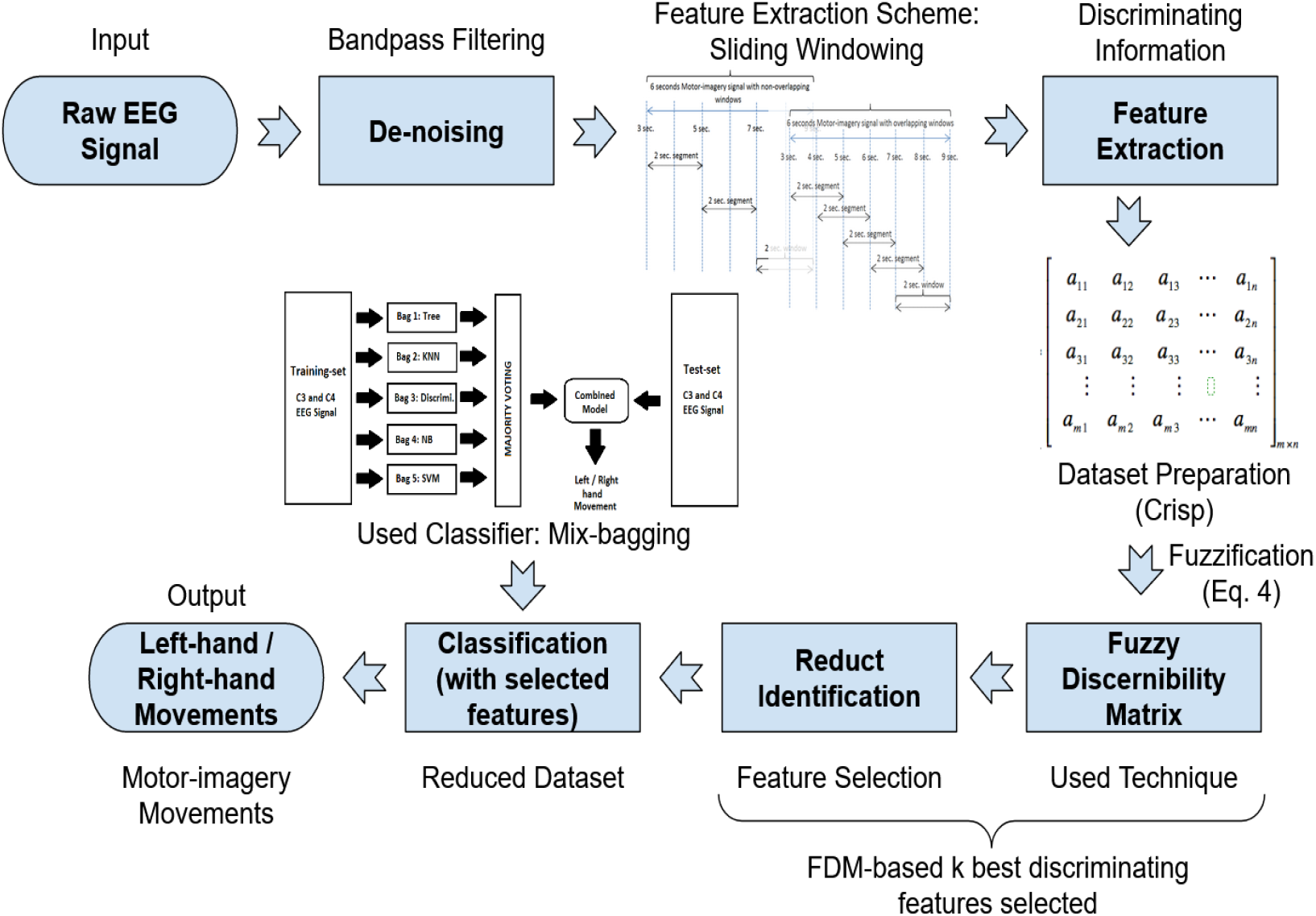
The diagram of the proposed end-to-end framework for an efficient EEG motor-imagery signal classification

## 4. Experimental Set-up

### 4.1. Dataset

The dataset contains raw EEG signals for three EEG electrodes *C*3, *Cz* and *C*4. The left-hand and right-hand movements are dominantly related to the brain regions tapped by the *C*3 and *C*4 electrodes. The region of our interest in each EEG signal starts at *t* = 3 seconds, lasts for 6 seconds and ends at *t* = 9 seconds. The IEEE 10 – 20 electrode placement and the signal description are shown in the Figures 7 and 8. This dataset does not have any major artifacts. However, the raw EEG signal is filtered with an elliptic band-pass filter using the cut-off frequencies 0.5 Hz and 50 Hz at a sampling rate of 128 Hz. The dataset has been used in this paper instead of other BCI standard datasets as it has longer available motor-imagery duration for each trial and therefore, is more suitable for the sliding window approach.

**Figure 7:**
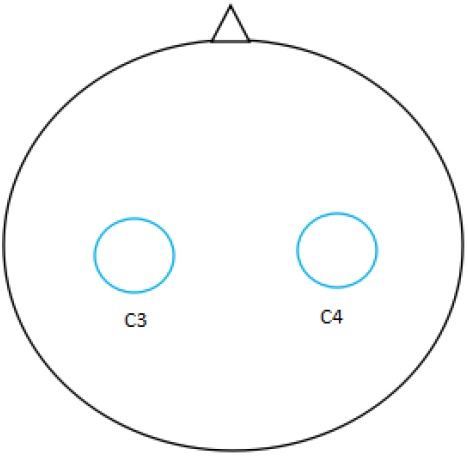
*C*3 and *C*4 electrodes placement based on IEEE 10 – 20 standard

**Figure 8:**
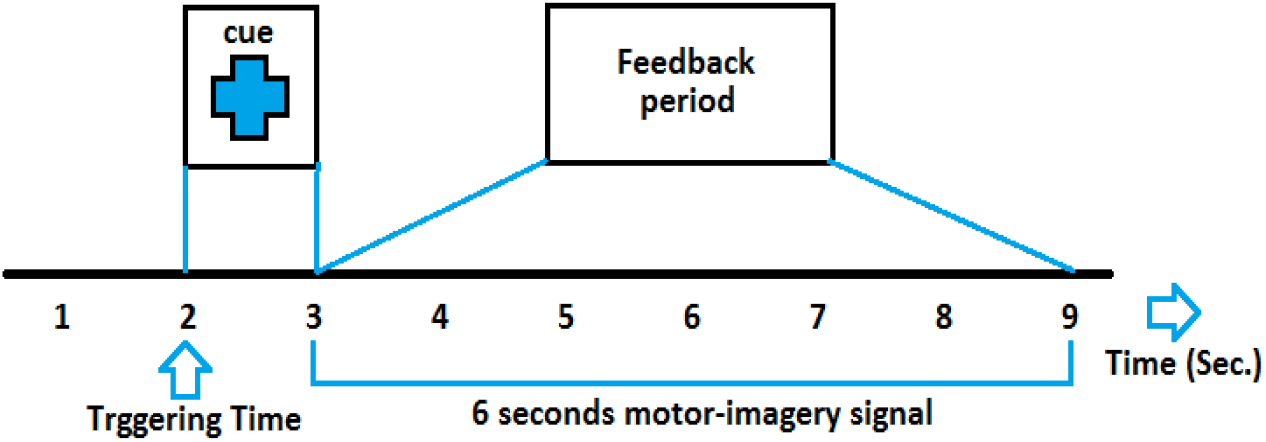
Raw EEG signal of 6 seconds long motor-imagery activity

The total dataset is of 280 trials (that is, samples) containing an equal number of the left hand and right-hand movement trials. The first 140 trials are taken for training-set and the remaining 140 trials are considered as test-set. To de-noise the input EEG signal, an elliptic band-pass filter with cut-off frequencies of 0.5 Hz and 30 Hz, has been employed. [14, 19, 20]. For easy understanding, the dataset description is elaborated in Table 6.

**Table 6:** Dataset Description.

| Dataset | Electrodes | Sample Size | Class Label | Training/ Testing | Cut-off Frequency |
| --- | --- | --- | --- | --- | --- |
| BCI competition II (dataset III) | C3 and C4 | 768 (6 sec. × 128Hz) | 1-left hand, 2-right hand movement | 140/140 | 0.5 – 30 Hz |

### 4.2. Experimental Framework

The used raw EEG signal is of 9 seconds. However, the signal extracted from the last 6 seconds is considered for our signal classification task. This is because the imagination of hand movements (right hand or left hand) is based on the initial 3 seconds cue and is recorded in the following 6 seconds. There are two 6 seconds signals recorded from two electrodes *C*3 and *C*4. Meaningful features need to be extracted from the complete 12 seconds signal. As of now, this 6 seconds trial is used to extract features considering it as a *single entity*. In this paper, DWT energy and entropy-based feature extraction has *not* been employed on the complete 6 seconds signal/per electrode; rather *multiple windows (each of* 2 *seconds) have been used to apply the energy entropy-based feature extraction for alpha (*8 – 13 *Hz) and beta (*13 – 25 *Hz) frequency bands separately*. It is done so because the imagination of the subject may vary in the 6 seconds signal and thus, a traditional feature extraction approach that considers the whole signal as a single unit may not capture detailed discriminative information. On the other hand, if the features are extracted from multiple overlapping (Fig. 9) or non-overlapping (Fig. 10) sliding segments of the original 6 seconds EEG signal based on alpha and beta EEG bands, they might provide us with additional information which is not captured in earlier cases [58, 59]. The window size has been kept at 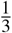 of the original signal length in our experiments. The number of features obtained from experiments Exp-(a), Exp-(b), and Exp-(c) are 8, 24, and 40 respectively.

**Figure 9:**
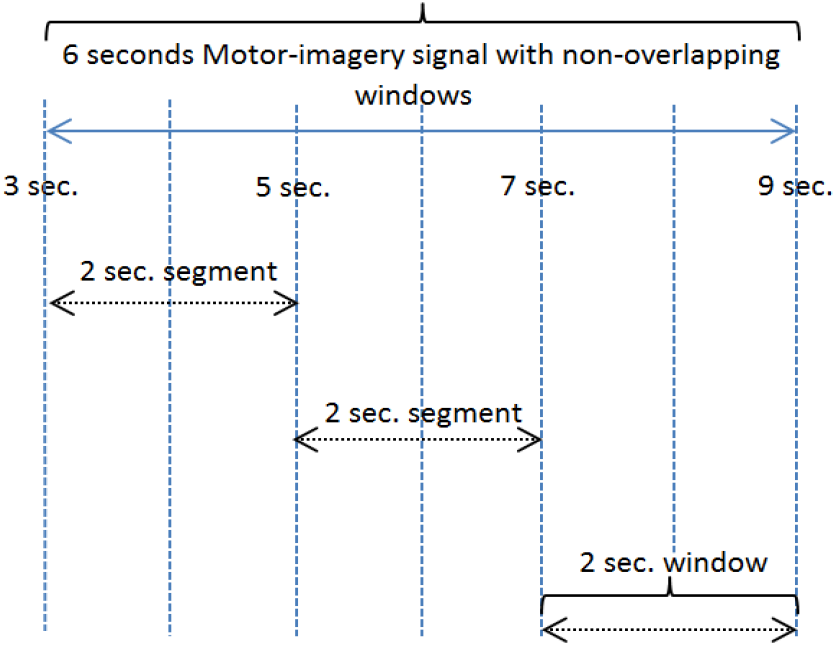
Diagrammatic representation of non-overlapping temporal window approach in Exp-(b)/Exp-(e).

**Figure 10:**
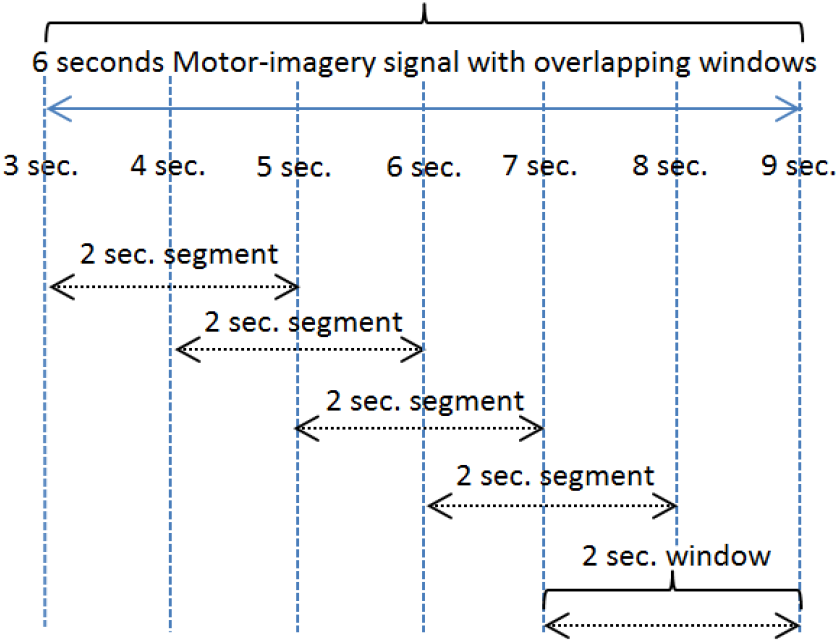
Diagrammatic representation of overlapping temporal window approach in Exp-(c)/Exp-(f).

Similarly in the tradition scheme of AAR, each electrode splits into alpha and beta frequency bands. AAR feature extraction technique with order=6 has been applied on the each sub-bands. It generates 12 features per electrode. Again, it is repeated for non-overlapping (3 segments) or overlapping (5 segments) feature extraction scheme. Subsequently, the process gives us 72 and 120 feature-sets respectively.

Therefore, we introduce a framework to extract, select and classify the most discriminating features from a EEG motor-imagery signal. DWT-based wavelet coefficients are used as primary features. Then, energy and entropy features are obtained from the features to consolidate the discriminating information and to reduce the primary feature-set length. Here, a sliding window strategy has been used to capture more embedded discriminating left-hand right-hand brain-states. There are three primary schemes for feature extraction explained in Table 7.

**Table 7:** Overview of our experiments.

| Experiment | Description |
| --- | --- |
| **Exp-(a)** | The wavelet energy and entropy features are extracted from Alpha and Beta sub-bands of the entire EEG signal (6 sec.) for each trial. |
| **Exp-(b)** | The whole EEG signal (6 sec.) for each trial is divided into three non-overlapping (2 sec. each, that is, 256 samples) segments as input and wavelet energy and entropy features are extracted from its Alpha and Beta sub-bands from each segment. (Figure 4) |
| **Exp-(c)** | The whole EEG signal (6 sec.) for each trial is divided into five overlapping (2 Sec. each, that is, 256 samples with 128 overlapping samples) segments as input and wavelet energy and entropy features are extracted from its Alpha and Beta sub-bands from each segment. (Figure 5) |
| **Note:** *Elliptic bandpass filter is used to extract different frequency bands from the EEG input signal* |  |

Once the features are obtained, a modified bagging ensemble classifier is used to classify them into left-hand/right-hand movement.

Our intuition before the experiments is that the use of overlapping or non-overlapping feature extraction technique, fuzzy discernibility matrix and modified bagging classification approaches can perform better over the traditional approaches. To analyze and validate our hypothesis, our study has been primarily segregated into two types of experiments: Experiment-I and Experiment-II.

The overlapping energy entropy approach also has been compared with AAR-based overlapping and non-overlapping approaches to re-examine its superior performance. The details of the AAR approaches with their comparison based on classification accuracy have been given in Tables 8 respectively.

**Table 8:** Overview of other experiments for feature-set comparison.

| Experiment | Description |
| --- | --- |
| **Exp-(d)** | The AAR (with order $p=6$) features are extracted from the entire EEG signal (6 sec.) for each trial. ⟨AAR normal features⟩ |
| **Exp-(e)** | The AAR (with order $p=6$) features are extracted from the non-overlapping (2 sec.) segments of the input EEG signal (6 sec.) for each trial. ⟨AAR non-overlapping features⟩ |
| **Exp-(f)** | The AAR (with order $p=6$) features are extracted from the overlapping (2 sec.) segments of the input EEG signal (6 sec.) for each trial. ⟨AAR overlapping features⟩ |
| **Note:** *Elliptic bandpass filter is used to extract different frequency bands from the EEG input signal* |  |

### 4.3. System Configuration

The experiments are implemented using MATLAB 2016a on an Intel(R) Core(TM) *i*5 – 6200*U* CPU 2.40 GHz with 64 bits Windows 10 Professional operating system and 8 GB RAM. The feature extractions, selection, and classifications of EEG signal are done by three separate MATLAB (.`m`) scripts. The feature selection based on the fuzzy discernibility matrix and mix-bagging classifier is coded in MATLAB. However, the rest of the used functions are MATLAB in-built methods tuned as per our parameter configuration.

## 5. Results and Analysis

### 5.1. Experiment-I

Two types of datasets have used (the AAR 12 and 24 features datasets) in Experiment-I. First, it compares the performances between the traditional discernibility matrix and our proposed fuzzy discernibility matrix-based feature selection techniques using hold out split (that is, first 140 trials for training-set and remaining 140 trials as test-set) using AAR-12 features dataset. Secondly, the proposed framework has been implemented on AAR-24 features dataset using same hold-out configuration.

The accuracy-based comparison between the traditional and the fuzzy discernibility matrix approaches along with their best discriminating features are given in Tables 9 and 10. We observe that the proposed fuzzy discernibility matrix with the dissimilarity measure defined in Eq. 6 performs best among the explored alternatives. The performance obtained from the other variants is not discussed while discussing the holdout technique. The proposed FDM (using Eq. 6) gives better accuracy not only to the traditional discernibility matrix-based feature selection technique but also the accuracy obtained using all the features without any selection technique. Although an exception is observed in AAR 24 features dataset when the *SVM*1 classifier applied on the FDM algorithm selected features subset, that is, accuracy 76.43%, it performs better than traditional technique but fails to outsmart accuracy 77.86% obtained while using all the 24 features. The second observation is that our proposed Mix-bag classifier consistently provides the best results for all our experimental cases in the hold out technique. In addition to the said observations, it is also found that the FDM and Mix-bag classifier combination gives us even better classification accuracy than the case where all the features used for classification. The classification accuracies obtained from AAR 12 and 24 feature datasets using the proposed Mix-bag classifier are 82.14% and 80.71%, respectively. Here, the user input (k) has been kept at 50% of the actual feature-set size.

**Table 9:** Results of accuracies (%) and selected features obtained from AAR 12 features dataset using holdout technique.

| Approach | SVM1 | SVM2 | ENS1 | ENS2 | Mix-bag | Selected Features | Feature-set Length |
| --- | --- | --- | --- | --- | --- | --- | --- |
| ALL Features | 79.29 | 78.57 | 77.13/3 | 78.57/6 | 81.43 | - | 12 |
| DM | 77.86 | 78.57 | 78.57/10 | 77.14/6 | 81.43 | 1,2,4,5,7,8,10,11,12 | 9 |
| FDM (Eq. 6) | 80.00 | 79.29 | 78.57/9 | 78.57/6 | **82.14** | 3,6,7,8,9,12 | 6 |

**Table 10:** Results of accuracies (%) and selected features obtained from AAR 24 features dataset using holdout technique.

| Approach | SVM1 | SVM2 | ENS1 | ENS2 | Mix-bag | Selected Features | Feature-set Length |
| --- | --- | --- | --- | --- | --- | --- | --- |
| ALL Features | 77.86 | 77.86 | 70.71/29 | 73.57/9 | 78.57 | - | 24 |
| DM | 62.14 | 63.57 | 56.43/3 | 62.86/6 | 65.71 | 7,10,20,22,23 | 5 |
| FDM (Eq. 6) | 76.43 | 78.57 | 71.43/6 | 73.57/6 | **80.71** | 3,11,12,13,14,15,18,19,20,22,23,24 | 12 |

These results indicate that the obtained selected feature subsets using FDM (with Eq. 6) have *higher discriminating information even over the original feature-sets*. In simple words, our experiments validate the notion that the use of all the features for a given dataset does not always ensure efficient classification due to low discriminating information of superfluous and redundant features. Therefore, feature selection is an indispensable part of a supervised learning paradigm. Thus our used Fuzzy Discernibility Matrix-based feature selection technique satisfies the purpose of enhancing the classification using the same feature extraction technique. Here, it must be mentioned that the reduct (feature subset) length is automatically computed in the traditional discernibility matrix-based feature selection. As it gives poor classification result with the reduct length 5 in Table 10, FDM with “k=5” has also been examined. It provide 76.43 % classification accuracy.

### 5.2. Experiment-II

The comparative performance of DWT-based energy entropy approaches (described in Table 7) with AAR-based approaches (described in Table 8) are given in Table 11.

**Table 11:** Comparative results obtained from different experiments along with their feature-set size using mix-bagging classifier.

| Experiment | Accuracy (%) | Feature-set size |
| --- | --- | --- |
| **Exp-(a)** | 84.29 | 8 |
| **Exp-(b)** | 88.57 | 24 |
| **Exp-(c)** | **91.43** | 40 |
| **Exp-(d)** | 81.43 | 24 |
| **Exp-(e)** | 84.29 | 72 |
| **Exp-(f)** | 89.29 | 120 |

There is a distinction between the results obtained from AAR-24 features (refer Table 10 where order of the AAR parameters is 12) and AAR traditional scheme of features obtained using alpha and beta sub-bands (that is, Exp-(d) and also the size is 24). However, the best accuracy (91.43%) is obtained using 40 features extracted from the experiment (c) (that is, Exp-(c)) using the mix-bagging classifier. Their performances can be represented as follows (where *A ≻ B* suggest performance of scheme A is better than scheme B):

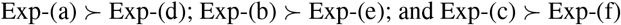

From the above comparison, it is concluded that the DWT-based energy entropy feature extraction schemes are superior in terms of discriminating information for a given input signal. Now, three different fundamental types of classifiers have been used with a total of eight variants. These classifiers are employed over datasets obtained from three distinct schemes. The hold-out technique is used for each model building and also executed the process several times independently to have a stable performance. The accuracies for each classifier is also calculated in Table 12 to get an overall impression of the best performing classifier. The top three best performing classifiers are mix-bagging, ENS3 and KNN. A comparison plot of the three best-performing classifiers has been given in Fig. 11 from our study in this paper.

**Table 12:** Accuracies (%) obtained in experiments with different classifiers using EngEnt.

| Experiments | NB | KNN | SVM1 | SVM2 | ENS1 | ENS2 | ENS3 | Mix-bagging |
| --- | --- | --- | --- | --- | --- | --- | --- | --- |
| Exp-(a) | 78.57 | 82.14 | 78.57 | 76.43 | 81.43 | 82.14 | 82.14 | 84.29 |
| Exp-(b) | 81.43 | 83.57 | 84.29 | 79.29 | 84.29 | 83.57 | 84.29 | 88.57 |
| Exp-(c) | 83.57 | 86.43 | 85.00 | 80.00 | 86.43 | 86.43 | 85.71 | **91.43** |
| Mean % | 81.19 | 84.05 | 82.62 | 78.57 | 84.05 | 84.05 | 84.05 | 88.10 |

**Figure 11:**
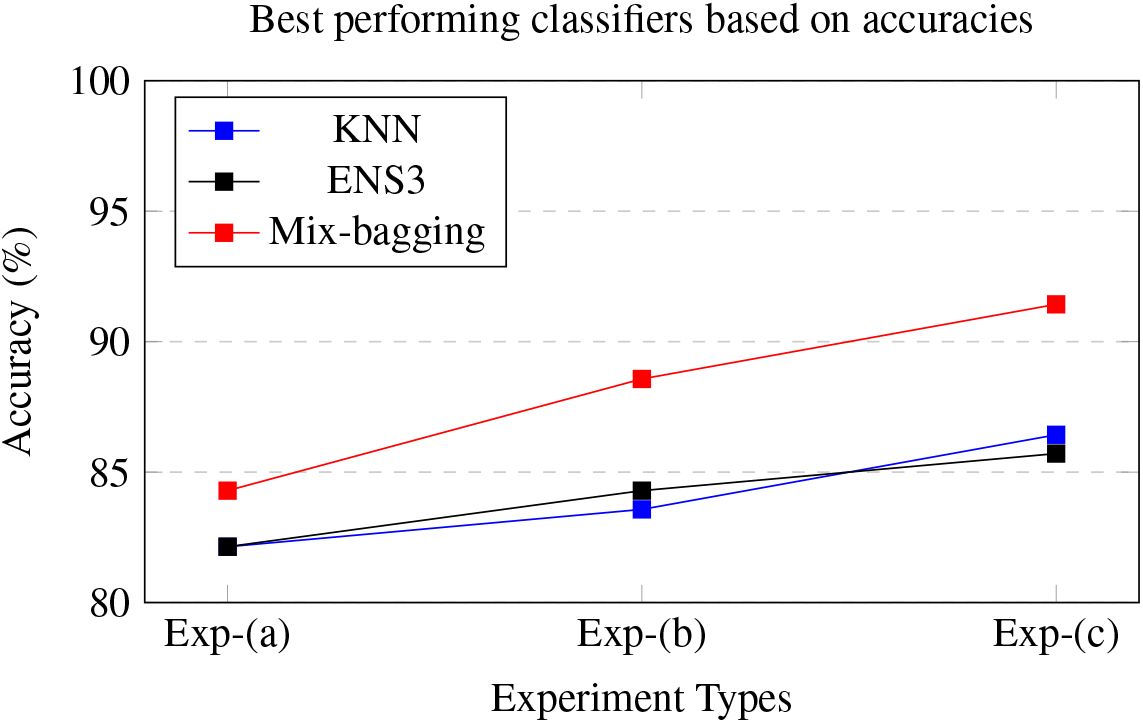
Plots of best performing accuracies obtained from different experiments using the actual feature-sets (without FDM)

As it has been already anticipated, overlapping and non-overlapping feature extractions schemes with DWT-based energy and entropy provide higher classification accuracies compared to the traditional feature extraction technique (that is, Exp-(a)). Eventually, the overlapping approach gives the best performance over the two other approaches mentioned in this paper. The comparative performance of all the three schemes can be represented as follows:

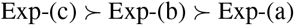

On the other hand, the mix-bagging classifier stands above all other classifiers due to the inherent diversity introduced by multiple types of learners. It exhibits better performance due to the advantages of all the learners while reducing their shortcomings through the ensemble. The proposed Exp-(c) using a mix-bagging classifier gives us the highest classification accuracy **91.43%**.

To downsize the used feature-set length, the FDM-based feature selection technique has been used [34, 46]. It is implemented with 25%, 50%, 75% features of the original feature-set length based on the discriminating power of the features. Performance obtained using different selected feature-subsets is shown in Table 13. It is observed that the Exp-(c) plus FDM-based feature selection technique with only 10 best discriminating features (that is, 25% of the actual feature-set length) gives us **92.14%**classification accuracy. This result is even better than the previous accuracy of 91.43% obtained using *all* the original features in Exp-(c). Thus, it not only reduces the feature-set size from 40 to 10 but also finds the most discriminant features for classification.

**Table 13:**
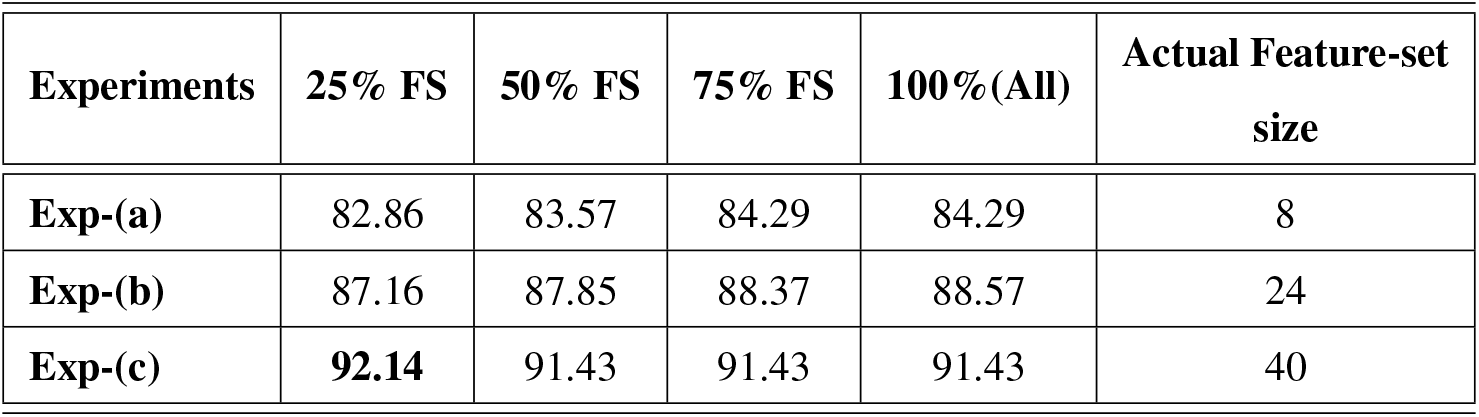
Accuracies (%) obtained from experiments based on different feature-set length (**FS**) using FDM.

| Experiments | 25% FS | 50% FS | 75% FS | 100% (All) | Actual Feature-set size |
| --- | --- | --- | --- | --- | --- |
| Exp-(a) | 82.86 | 83.57 | 84.29 | 84.29 | 8 |
| Exp-(b) | 87.16 | 87.85 | 88.37 | 88.57 | 24 |
| Exp-(c) | **92.14** | 91.43 | 91.43 | 91.43 | 40 |

Our top two proposed approaches provide the best-known classification accuracy for the used BCI dataset. Finally, a comparison of various existing best techniques and the proposed method is given in Table 14. Fig. 12 graphically compares the accuracy produced by the best two approaches in this paper with other classification methods applied to this dataset to date.

**Table 14:** List of existing few best performing techniques on BCI Competition 2003 dataset III.

| Reference | Method used | Classifier | Accuracy (%) |
| --- | --- | --- | --- |
| Chatterjee et al. [20] (Chatt1) | Wavelet Energy-entropy | SVM (kernel: Linear / polynomial) | 85.00 |
| Chatterjee et al. [19] (Chatt2) | Average-power + band -power + Wavelet Energy-entropy + RMS + Statistical features (Table V) | MLP | 85.71 |
| Proposed1 Exp-(c) | Overlapping temporal window + Alpha-beta bands + Energy-entropy | Mix-bagging ensemble | 89.29 |
| Lemm et al. [21] | Morlet wavelet | Bayes Quadratic | 89.29 |
| Bashar et al. [22] | MEMD + STFT | KNN (cosine distance) | 90.71 |
| Proposed1 Exp-(c) | Overlapping temporal window + Alpha-beta bands + Energy-entropy | Mix-bagging ensemble | 91.43 |
| Proposed2 Exp-(c) | Overlapping temporal window + Alpha-beta bands + Energy-entropy + FDM (Eq. 6) | Mix-bagging ensemble | **92.14** |

**Figure 12:**
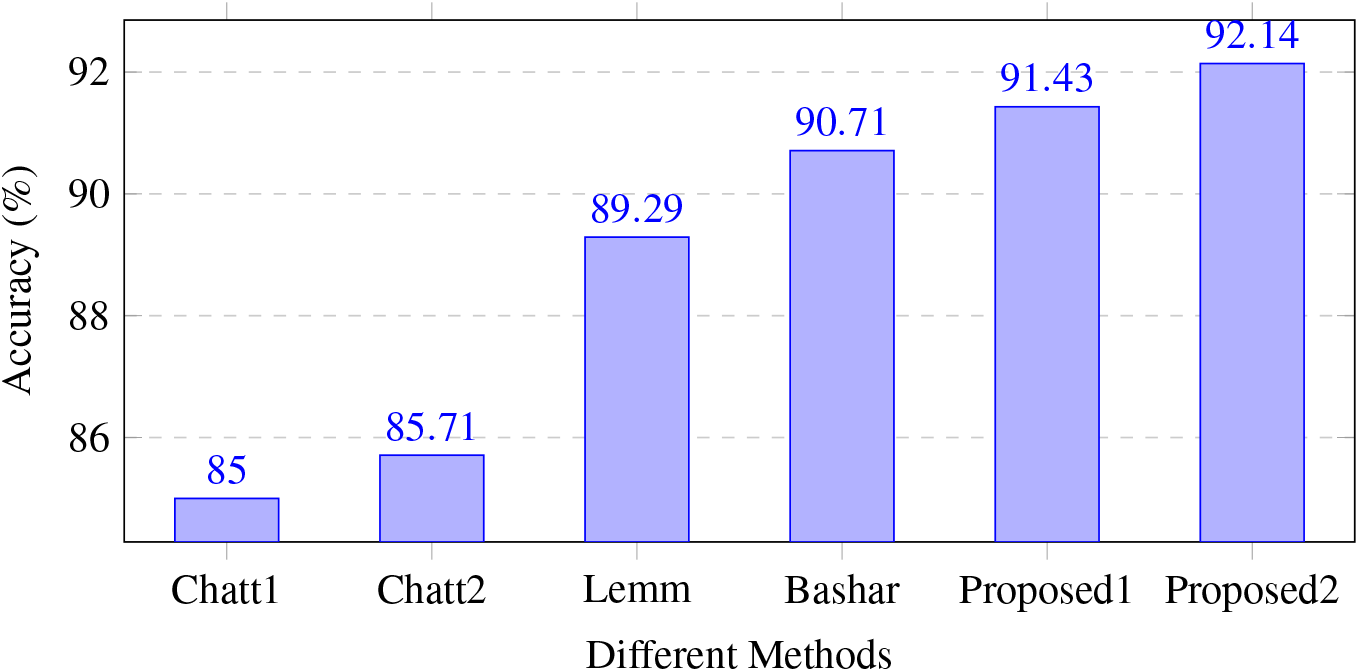
Accuracies of a few best performing techniques on BCI Competition II Dataset III.

We observe that the results obtained from our proposed approaches are truly aligned with our initial anticipation that the sliding window with wavelet energy and entropy feature extraction technique provides more insights in terms of *discriminative information* due to the use of multiple temporal segments than that provided by the traditional approach on the same trial.

## 6. Conclusion

Motor-imagery EEG signals and their classification is the prime objective of this paper. A new framework of feature extraction, selection, and classification have been used so that the cognitive task encoded in the EEG signal can be automatically inferred. It is significant in the design and deployment of future BCI devices. Cognitive tasks that involve physical muscle movements or even the imagination of different limb movements develop similar signatures in brain activities. In EEG-based motor-imagery signal classification, the primary aim is to capture the unique patterns associated with each brain state efficiently. As the EEG signals are non-stationary, the feature extraction technique which extracts both the time domain and the frequency domain information from the motor-imagery EEG signal contains the most discriminating information of the left-hand and right-hand movements. The segment of the signal where the exact discriminating information resides is not fixed even in all the the trials of the same subject. Therefore, the extraction of features from the exact region of activities improves the classification performance. The used feature extraction technique induced with the established discrete wavelet transform energy and entropy provides the relevant insights of both the spectral and temporal feature-space. However, the number of features increases as the number of overlapping windows goes up. More features do not necessarily guarantee good classification accuracy. In a highly correlated feature-set, all the features do not contribute in building an efficient classification model. Then, it is better to select the best discriminating features from the original feature-set. A suitable feature selection technique not only reduces the feature-set length but also increases the accuracy by discarding redundant non-discriminatory features.

The fuzzy discernibility matrix-based feature selection technique has been used to find the most discriminating feature-subset from the original feature-set, which in turn reduces the size of the selected feature-set. The combined proposed approaches of *feature extraction* and *classification* achieve a classification accuracy of 91.43%, which is higher than the reported best-ever performance on the BCI Competition II dataset III. We have further improved the classification accuracy to 92.14% when the proposed feature *selection* technique is implemented in addition to the remaining proposed techniques.

## Compliance with ethical standards

### Conflict of interest

The authors declare that they have no conflict of interest.

## Footnotes

1 We use the shorter notation to mean the fuzzy set 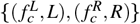

## Notes

### Competing Interest Statement

The authors have declared no competing interest.

### Summary of Updates

The title of the paper has been revised.

